# Informatics for Cancer Immunotherapy

**DOI:** 10.1101/152264

**Authors:** J. Hammerbacher, A. Snyder

**Affiliations:** Department of Genetics and Genomic Sciences, Icahn School of Medicine at Mount Sinai, New York, NY 10029; Department of Microbiology and Immunology, Medical University of South Carolina, Charleston, SC 29425; Department of Medicine, Memorial Sloan Kettering Cancer Center, New York, NY 10065; Department of Medicine, Weill Cornell Medical College, New York, NY 10065

**Keywords:** computational biology, bioinformatics, immunotherapy, neoantigens, checkpoint blockade, adoptive cell transfer

## Abstract

The rapid development of immunomodulatory cancer therapies has led to a concurrent increase in the application of informatics techniques to the analysis of tumors, the tumor microenvironment, and measures of systemic immunity. In this review, the use of tumors to gather genetic and expression data will first be explored. Next, techniques to assess tumor immunity are reviewed, including HLA status, predicted neoantigens, immune microenvironment deconvolution and T-cell receptor (TCR) sequencing. Attempts to integrate these data are in early stages of development and are discussed next. Finally, we review the application of these informatics strategies to therapy development, with a focus on vaccines, adoptive cell transfer, and checkpoint blockade therapies.

## Introduction

The quest to understand and improve how the immune system identifies and eradicates cancer can make use of data from dozens of molecular, cellular, and tissue profiling technologies. The primary focus of this review will be bioinformatic analyses of data generated by high-throughput DNA and RNA sequencing of bulk normal and tumor cell populations.

First, computational tools that consider cancer and the immune system separately will be discussed, followed by a discussion of how to better understand the interaction of the immune system and cancer. Finally, methods to dissect therapeutic responses to interventions such as vaccination, checkpoint blockade, and adoptive cell transfer will be reviewed.

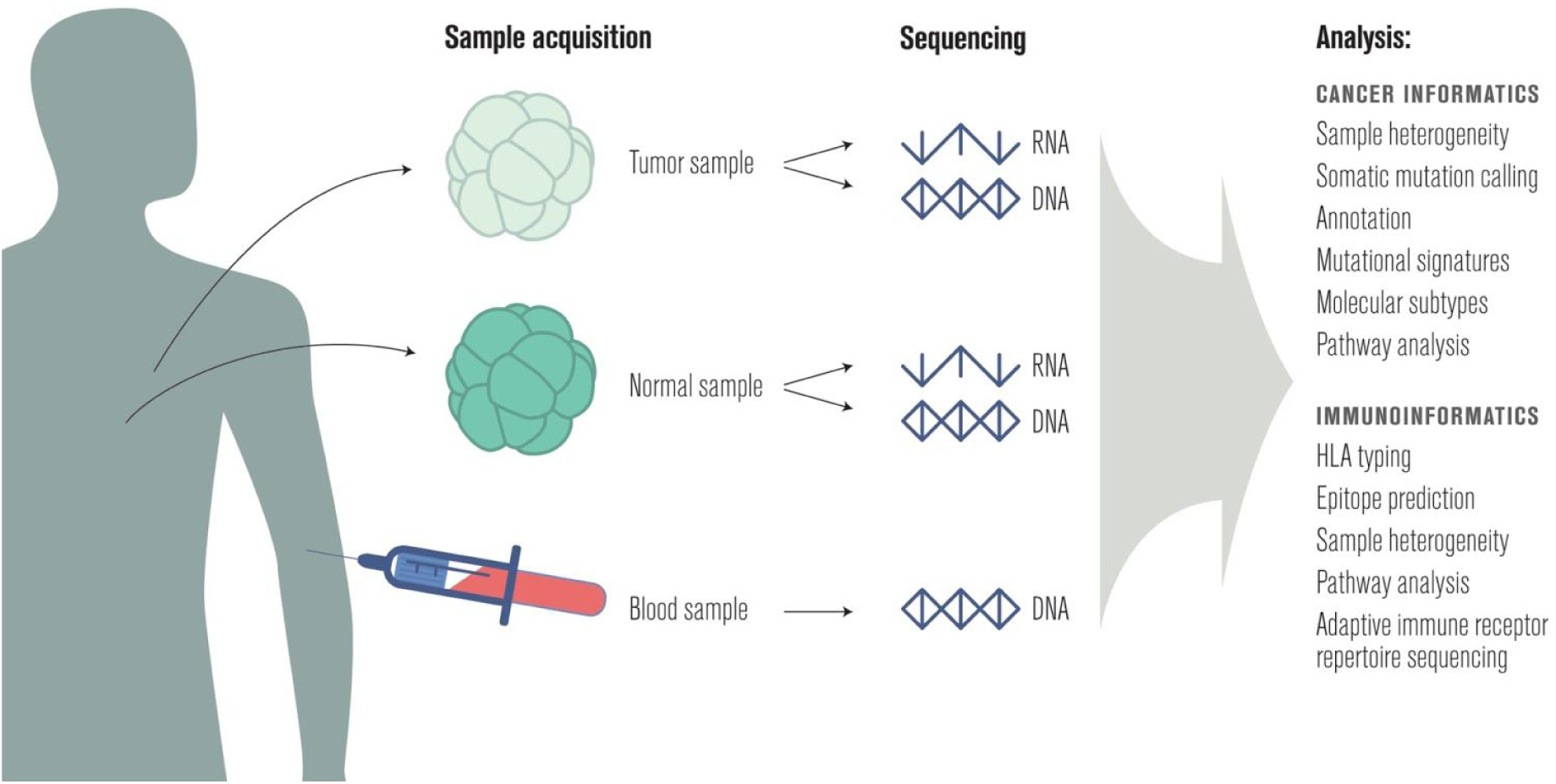

## Cancer informatics

Cancer is a disease of the genome [1], so it is common to profile the mutations and transcripts present in both the patient’s peripheral blood or nearby normal tissue and one or more tumor samples [2]. A summary of common uses of this data in cancer research follows; for a more detailed review see [3].

### Sample heterogeneity

The proportion of tumor cells in a cancer tissue sample is known as the “cellularity” or “purity” of the sample. Cellularity is an important metric for any type of downstream analysis since it is directly associated with the intensity of the tumor signal in genomic data. Inferring cellularity is complicated in some cancers because of challenges involved with pathologically excising normal tissue; in addition, nearby cells that appear normal microscopically often harbor somatic mutations similar to those found in cancer cells [4]. ABSOLUTE [5] and ESTIMATE [6] are two commonly-used tools to compute cellularity from sequencing data; recent work compared these tools to each other and to pathology-based estimates of cellularity and found low, but positive, correlation between sequence-based and pathology-based estimates [7]. While ESTIMATE uses gene expression as input, ABSOLUTE uses somatic copy number to quantify cellularity and hence is sensitive to errors in copy number estimates.

Within the population of tumor cells there is heterogeneity [8, 9]. Tumor cells can be partitioned into clonal families, and it is sometimes possible to reconstruct the phylogeny of tumor cells to better understand the spatial and temporal evolution of tumor heterogeneity [10–14]. Tools like THetA [15], PyClone [16], SciClone [17], PhyloWGS [18], QuantumClone [19], and Canopy [20] have been developed to compute tumor phylogenies, and visualization tools like BubbleTree [21] and fishplot [22] can be useful to better comprehend tumor evolution [23]. These tools operate on genomic data obtained from either a single tumor sample (THetA, PyClone, SciClone, Canopy, bubbletree) or multiple samples (QuantumClone, fishplot) that are spatially or temporally distinct from each other.

Recently, single-cell sequencing technologies have been employed to improve sample heterogeneity estimates [24–26]; these approaches are reviewed in [27].

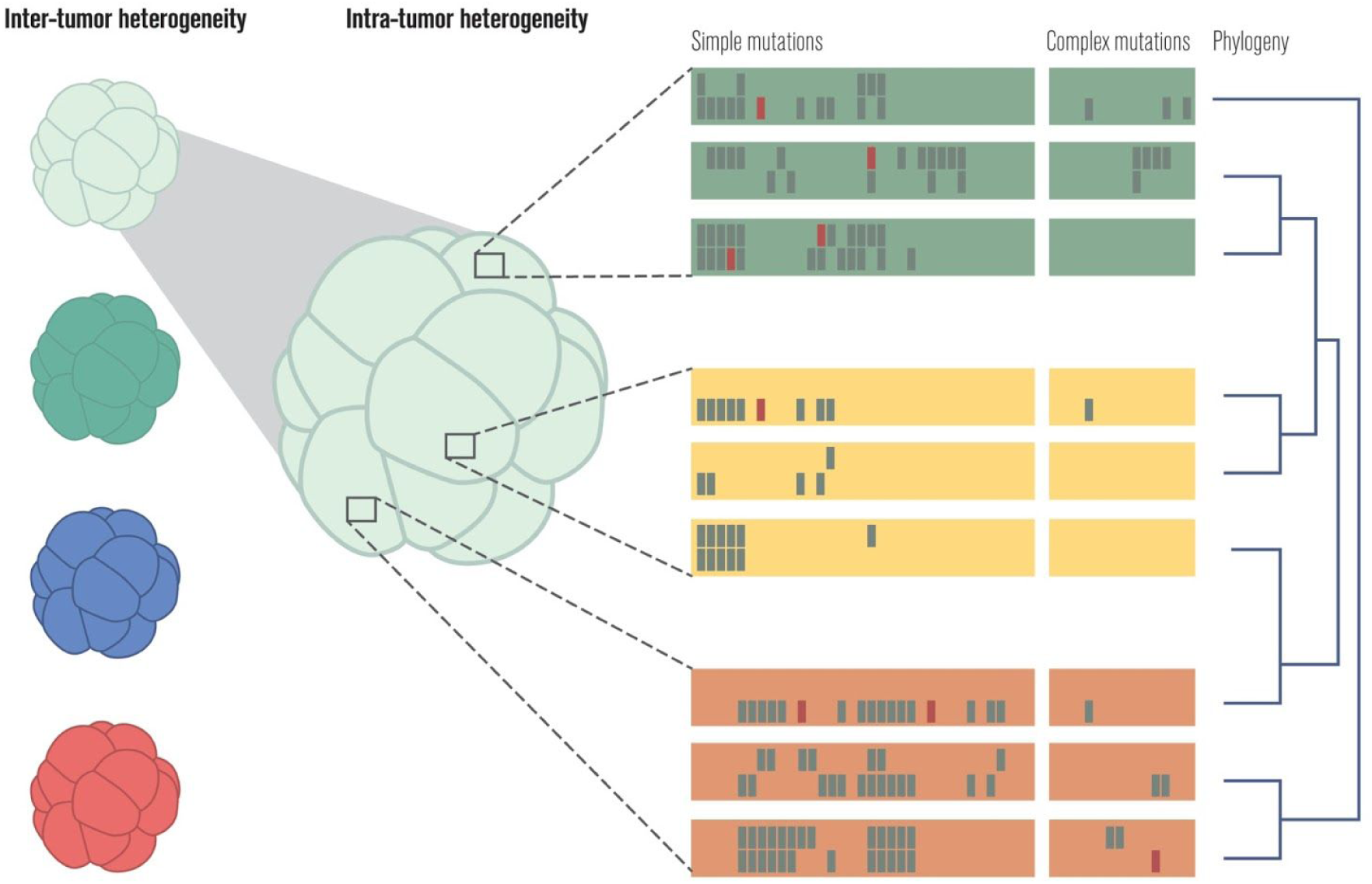

### Somatic mutation calling

The Genomic Data Commons [28] makes available single nucleotide variant and short insertion and deletion (indel) calls from MuTect2, SomaticSniper [29], VarScan2 [30], and MuSE [31]. A recent ICGC benchmarking exercise [32] made use of two additional callers, MuTect [33] and Strelka [34]. In practice, it is common to combine the results of several somatic mutation callers to produce a consensus list of high-confidence calls [35, 36].

While simple mutations are the most frequent form of genomic alteration in cancer cells, more complex mutations are almost always present and are often oncogenic drivers. Complex mutations can be organized into categories [37], and it is common to use a different tool for each category of complex mutation.

Software for structural variant detection includes DELLY [38], Meerkat [39], and novoBreak [40]. It is helpful to have whole genome DNA-seq data when using these tools, since these variations often affect large chromosomal regions, and are difficult to detect with whole exome DNA-seq data.

Two recent benchmarks [41, 42] of gene fusion detection software found that EricScript [43], SOAPfuse [44], FusionCatcher [45], and JAFFA [46] performed well. These tools make use of RNA-seq data as input.

Somatic copy number alterations (SCNAs) for the TCGA project were estimated with ABSOLUTE [47]. Recent benchmarks [48, 49] of software for SCNA calling from sequencing data highlighted ADTEx [50] and EXCAVATOR [51] as top performers. Tools not included in those benchmarks that may be useful include seqCNA [52], Sequenza [53], hapLOHseq [54], and CNVkit [55].

Challenges to the field of bioinformatics include the need for continuous updates to these programs, and the conflict between repeated benchmarking to delineate the best program versus consistent use of the same program to compare results between studies.

The somatic mutation callers discussed above make assumptions about sequencing protocols and read preprocessing software and methods. For example, many simple mutation detection methods expect whole exome data, and the choice of the exome capture kit can alter the somatic mutations identified. Sequencing depth is a critical consideration for detecting somatic mutations, as higher depth can better resolve sample heterogeneity issues. Usually one normal and one tumor genome are sequenced; however, some somatic mutation callers try to infer somatic mutations without a normal sample [56], while others can make use of RNA-seq data [57, 58] or data from DNA-seq of multiple samples. Biopsy and tissue preservation methods can also impact somatic mutations detected [59, 60].

Standard read preprocessing includes alignment to the human reference, base quality score recalibration, duplicate removal, and realignment of indels [61]. It is also important to profile the quality of the sequencing data and the alignments [62]. Significant coordination of many software tools is required to implement quality control, alignment, post-alignment processing, and both simple and complex mutation calling [63]. Most labs still manually examine somatic mutation calls with tools like IGV [64] or pileup.js [65] to confirm their accuracy, as in [32]. Whole pipelines have been published that can serve as a model for investigators setting up their own computational programs [66–68]. As of 2017, there is no consensus on the most sensitive and specific pipeline.

In fact, despite years of work on somatic mutation calling, it is still common for two labs to produce highly discordant results on the same input material, even for simple mutations called from targeted panels [69] or whole exome sequencing [70]. Attempts to benchmark somatic mutation calling pipelines often consider only concordance comparisons [71] or validate on simulated data [72], and the most commonly used validation technique, Sanger sequencing, is time consuming and not always accurate [73]. Recently, in an effort to address this quandary and provide model data, high-coverage whole genome, whole exome, and targeted panel data have been made available for a single AML case [74]. While this high coverage data will be useful, it still does not capture the significant impact of variation in sequencing protocols across sites [75].

Sample heterogeneity further complicates the validation process. Ultimately, a standards body should provide isogenic cell lines that can allow for benchmarking of a site’s somatic mutation calling process, similar to Genome in a Bottle for germline mutation calling [76].

### Annotation

After enumerating high-confidence somatic mutations in a tumor sample, the next step is to annotate each mutation with its expected impact. Basic annotations include whether the mutation occurs within a gene, whether it alters the protein produced from that gene, and the specific amino acid alterations expected. Cancer-related annotations include whether the mutation occurs within a “driver” gene [57] that is known to contribute to the progression of cancer and whether a drug exists that can target cells carrying the mutation.

Common tools for the annotation of somatic mutations include VEP [77], SnpEff [78], ANNOVAR [79, 80], and Oncotator [81]. These tools make use of databases like ENSEMBL [82, 83], RefSeq [84], and the UCSC Known Genes database [85] for basic annotations, and COSMIC [86], CIViC [87], and PMKB [88] for cancer-related annotations. The calling and annotation of somatic mutations is reviewed in [89]. It is important to note that there are differences among the annotation databases that can impact downstream analyses [90–92].

### Mutational signatures

Researchers recently disaggregated the biological processes generating somatic mutations in cancer by examining the trinucleotide context of somatic mutations in thousands of tumor samples [93]. These mutagenic processes include disruptions to the DNA damage repair pathway [94], environmental insults like smoking [95] and UV radiation, and chemotherapy [96]. Tools like deconstructSigs [97] facilitate the determination of the mutagenic processes present in a single tumor sample. Lessons from several years of mutational signature detection were recently distilled in a review [98].

### Molecular subtypes

In addition to the intra-tumor heterogeneity discussed above, tumors also vary across individuals [99] and can be categorized in a variety of ways, including clinically (e.g., by organ and stage) and pathologically (e.g., by grade and cell morphology). DNA-seq and RNA-seq data, the focus of this review, can also be used to categorize cancers into subtypes [100]. It is common to start with a single cancer, as categorized by organ or tissue of origin, and identify molecular subtypes using DNA-seq, RNA-seq, and other forms of sequencing data. Many of the TCGA research network publications attempt to identify molecular subtypes of a specific cancer in this way, for example in the cases of head and neck squamous cell [101] and clear cell renal cell [102] carcinomas. Another approach is to look for molecular subtypes across a variety of cancers [103].

### Pathway analysis

Systems biologists have mapped the hallmarks of cancer [104] and other cell behaviors onto the underlying protein-protein interaction and transcriptional regulatory networks that implement the behaviors [105–107]. These gene sets or pathways are collected in databases like KEGG [108, 109], GeneSetDB [110], MSigDB [111], and Reactome [112].

Grouping altered genes according to these gene sets and pathways increases the statistical power of computational analyses and also permits a higher-level view of altered cellular mechanisms based on our prior information on genes’ roles. Tools like Gene Set Enrichment Analysis (GSEA) [113, 114] and Ensemble of Gene Set Enrichment Analyses (EGSEA) [115] leverage gene sets for a better comparison of two groups of samples. This type of analysis is commonly used when comparing normal samples to tumors, tumors to a panel of normal samples (when the matched normal sample is not available) [116], or two sets of tumor samples that have different phenotypes (such as responders and non-responders). When a more granular understanding of the altered pathways for individual samples is needed, it is more common to use tools like single sample GSEA (ssGSEA) [117], gene set variation analysis (GSVA) [118], and moGSA [119] that perform single-sample gene set or pathway enrichment analysis.

## Immunoinformatics

As immune modulatory agents join the armamentarium of anti-cancer therapies, bioinformaticians involved in translational studies are increasingly using tools that profile the state of the immune system, both local and systemic.

### HLA typing

While the immune system has many cell types, the primary effector cells of tumor immunity are T cells [120]. T cells are activated in part by major histocompatibility complex (MHC) proteins presenting peptides for inspection on the surface of target cells [121]. In humans, the MHC genes are known as human leukocyte antigens (HLA), and they vary widely across individuals. There are three Class I and three Class II HLA genes, with the former recognized by CD8+ T cells and the latter by CD4+ T cells [122]. Enumerating the 12 alleles of these genes present in an individual is known as HLA typing. Most HLA typing methods make use of a database of known HLA alleles that is maintained by International ImMunoGeneTics (IMGT) [123].

High-resolution HLA typing can now be performed with DNA or RNA sequencing rather than serology [124]. Software for HLA typing includes ATHLATES [125] and POLYSOLVER [126], which take whole exome DNA-seq data; seq2HLA [127], which works with RNA-seq data; and OptiType [128], which works with either DNA-seq or RNA-seq data. A recent benchmark of seven algorithms found OptiType to be the most accurate on whole exome DNA-seq data for Class I HLA typing [129]. Promising new directions for HLA typing include using long read DNA-seq [130–132] and representing the polymorphic HLA region as a graph [133].

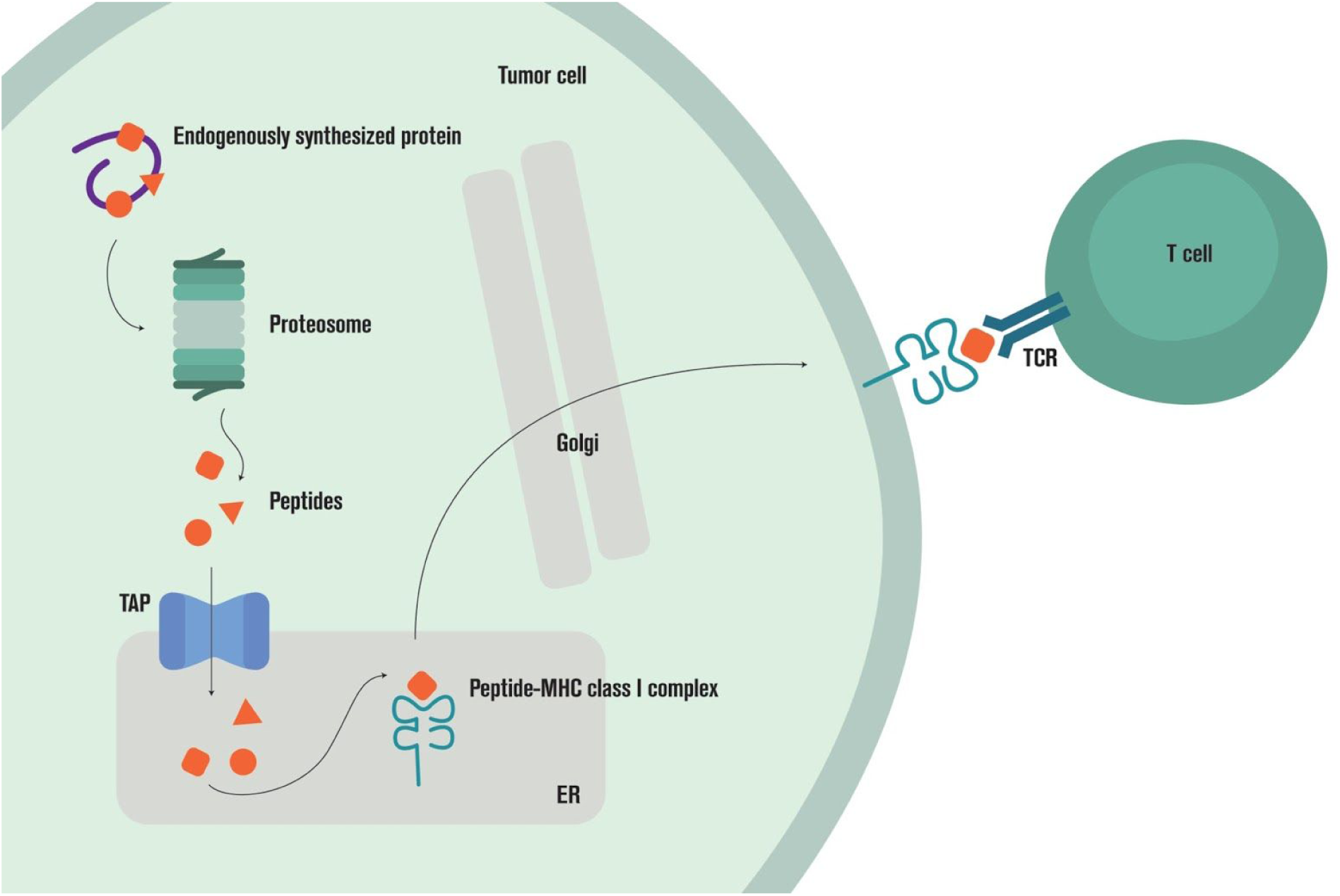

### Epitope prediction

The antigen processing machinery (APM) of a cell is responsible for sampling proteins and peptides from the cell interior for display to T cells [134]. If a presented peptide generates a T cell response, it is referred to as an epitope and said to be immunogenic.

There are many stages in the Class I APM pathway that can be predicted, including proteasomal cleavage, MHC binding, and presentation. The Immune Epitope Database (IEDB) [135] has curated measurements of each stage. Using IEDB data as input, researchers have built comprehensive predictors of how the Class I APM pathway will process a protein [136]. The most critical value to predict as a proxy for immunogenicity appears to be the binding affinity of peptide to the MHC protein [137]. Predictors for p/MHC binding affinity have been progressively improving for decades, led by the NetMHC family of predictors, accounting for complexities such as HLA alleles with little or no training data [138], variable peptide lengths [139], and variable binding affinity distributions across HLA alleles [140]. Other top performing predictors on the IEDB MHC Class I benchmark [141] include SMMPMBEC [142], MHCflurry [143], and the IEDB Consensus predictor [144].

Class II epitope prediction is currently far more difficult than Class I because the Class II APM pathway is complex [145], and Class II epitopes have more variable lengths and binding positions. For a recent review of Class I and Class II epitope prediction software, see [146].

Recently, advances in mass spectrometry have allowed direct and high-throughput measurement of presented peptides [147–150], so it is likely that presentation will soon replace MHC binding affinity as the most common outcome to predict as a proxy for immunogenicity. Beyond presentation, high-throughput measurements of TCR interactions with pMHC complexes are in development [151, 152]. It may one day be possible to predict pMHC/TCR interactions rather than p/MHC binding or pMHC presentation.

Predicting epitopes is complicated by our incomplete understanding of the origin of the peptides presented by a cell [153]. Some have suggested that defective ribosomal products (DRiPs) account for a significant fraction of presented peptides [154]; others claim that proteasome-generated spliced peptides are important [155–157]. There is even evidence that pMHC complexes can be transferred between cells [158]!

Also complicating epitope prediction are post-translational modifications [159, 160] and epitopes found on non-classical MHC proteins like HLA-E [161–163].

### Sample heterogeneity

With DNA and/or RNA sequencing data from bulk tissue, it is sometimes possible to estimate the relative proportion of immune cell types present in the sample. This computation is referred to as “deconvolution” [164, 165] and usually relies on the identification of “signature” genes whose expression levels can be used to distinguish between cell types [166]. Software tools for immune infiltrate deconvolution include CIBERSORT [167], TIMER [168], and MCP-counter [169].

These tools are often trained on expression data from isolated immune cell subtypes such as those generated by the Immunological Genome Project (ImmGen) [170] and differ in the number and kind of cell subtypes estimated: TIMER estimates 6 immune cell subtypes, MCP-counter estimates 8 immune cell subtypes and 2 stromal cell subtypes, and CIBERSORT estimates 22 immune cell subtypes. CIBERSORT and TIMER estimate relative frequencies of immune cell subtypes while MCP-counter can estimate absolute abundances. Users should take care when using CIBERSORT on RNA-seq data, as the model is trained on expression array data.

### Pathway analysis

Tools for gene set and pathway analysis can be used on populations of immune cells just as they can be on cancer cells. While gene sets and pathways for immune cell behaviors such as antigen processing, interferon response, and cytolysis are available from many of the same databases that hold data on cancer pathways, there are also immune-specific databases like DC-ATLAS [171] and InnateDB [172].

### Adaptive immune receptor repertoire sequencing

New library preparation, sequencing, and analysis technologies have enabled the high-throughput sequencing of B and T cell receptor repertoires [173–175]. The original protocols could only sequence unpaired single chains of these receptors; recent protocols make it possible to sequence paired chains [176, 177] to give a complete picture of the receptor repertoire. New algorithms allow repertoire sequences to be extracted from DNA-seq or RNA-seq data from bulk tissue [178–180] or single cells [181].

Public databases of repertoire sequences [182] allow individual samples to be placed into a larger context, and the community is actively working to standardize file formats and processing pipelines [183, 184].

Repertoire sequence populations can be summarized and compared across conditions to understand vaccination [185], aging [186], and more [187, 188]. Receptor diversity is an often-computed summary metric [189], and individual receptors are frequently grouped into clonal families as an intermediate step in these analyses [190]. The Repertoire Dissimilarity Index is another recently proposed measure that can be used to compare two samples [191].

## Cancer-immune interactions

Cancer and the immune system are both complex entities that interact and evolve over time. Milestones in the analysis of their interactions include work on cancer immunosurveillance [192–194], immunoediting [195–198], immune escape [199], and the cancer-immunity cycle [200].

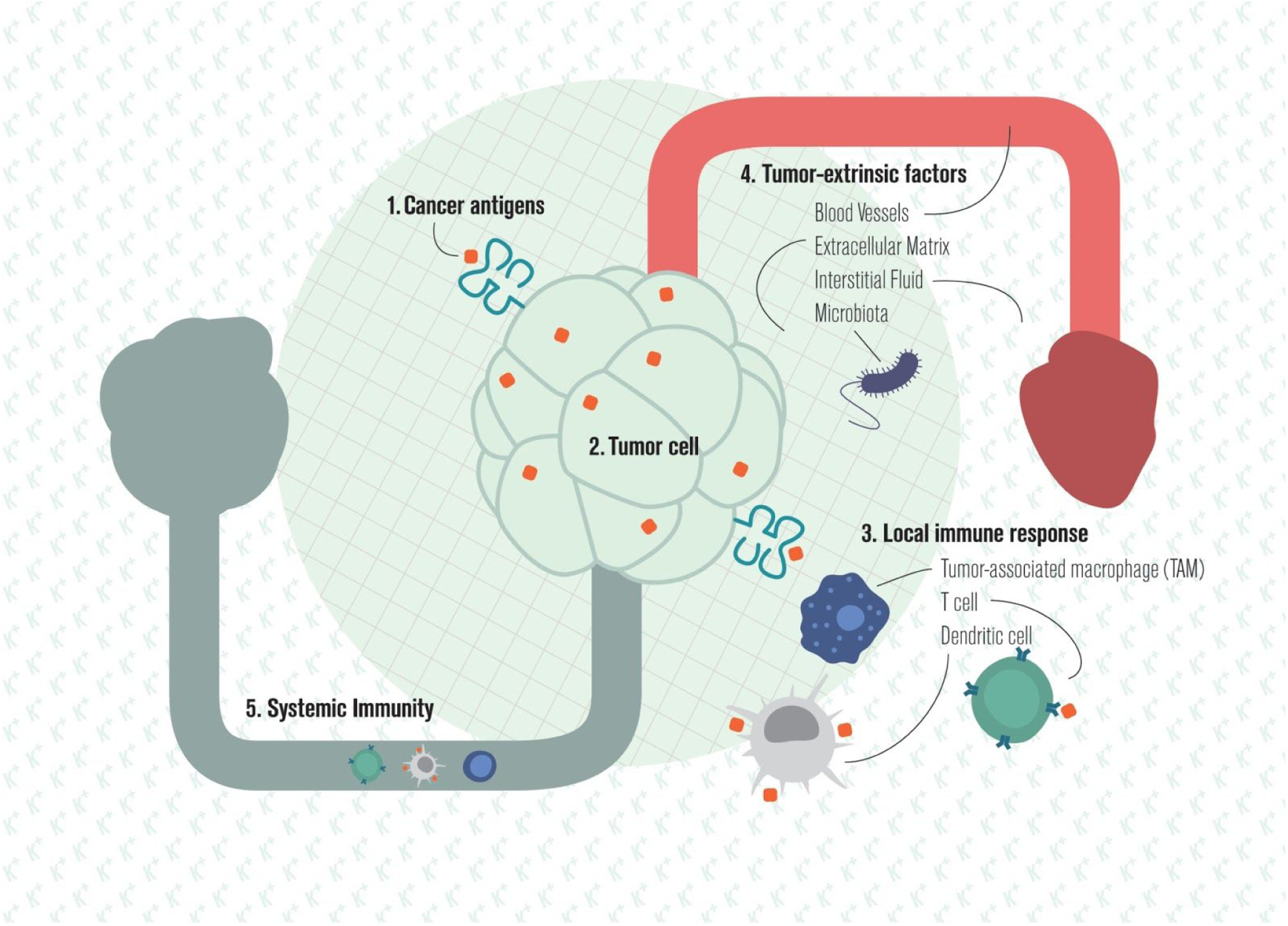

### Cancer antigens

What are the specific targets of the T cell response to cancer [201]? Many are self-antigens that are aberrantly expressed or post-translationally modified [202], while others are non-self viral [203] or somatic mutation-derived tumor antigens [204]. Somatic mutation-derived tumor antigens are also called “neoantigens” and are important targets for tumor rejection [205–210].

The feasibility of identifying neoantigens with whole exome DNA-seq data was demonstrated in 2008 [211] and has since been used to interrogate the neoantigen-directed response in many cancers including CLL [212]. Open source software for computing neoantigens from sequencing data includes Epidisco [213], ProTECT [214], and pVAC-Seq [215]. The TRON Cell Line Portal (TCLP) catalogs predicted neoantigens for common cancer cell lines [216]; The Cancer Immunome Atlas (TCIA) does the same for 20 solid tumor types from TCGA [217]. A recent analysis of predicted neoantigens for over 60,000 patients found very few shared neoantigens [218].

Recently it has become possible to directly identify peptides presented by cancer cells biopsied from human patients using mass spectrometry [219] and to screen for T cells specific for >1,000 peptide targets in a single tumor sample with high-throughput MHC tetramer assays [220]. Data from these assays is expected to improve the quality of cancer neoantigen prediction.

While the neoantigen burden of a tumor associates with patient survival [221], this effect cannot be separated from the impact of mutation burden [222]. The field is currently working to determine if specific mutagenic processes like chemotherapy are more likely to create neoantigens than others [96], and if certain kinds of neoantigens are more predictive of response than others. It is common to use RNA-seq data to lower the ranking of neoepitopes with no evidence of expression and to exclude neoepitopes determined to be similar to self-peptides. HLA binding affinity of validated neoepitopes and their corresponding unmutated wild type peptides have been closely examined with mixed results [223, 224]. Clonal neoantigens appear to be better than subclonal ones at eliciting a productive anti-tumor immune response [225].

### Tumor-intrinsic immune escape

As tumors grow, their constituent clones evolve to evade the immune system [226–229]. Some direct mechanisms of evasion include resistance to T cell lysis [230], or elimination of T cell targets through downregulation of the APM or editing out cancer antigens [134, 231]. Cancer cells also upregulate inhibitory checkpoint ligands such as PD-L1 through a variety of mechanisms [232–236]. While the impact of interferon on cancer cells is complex and still being elucidated, there is strong evidence that the disruption of interferon response pathways plays an important role in immune evasion [237–239].

Sometimes pathways whose primary function upon activation is to equip the tumor with a hallmark of cancer behavior have immune suppression as a secondary function, such as the MAPK [240, 241] and WNT [242] pathways. Recent evidence indicates that large chromosomal disruptions, common in cancer cells, may aid immune evasion [243].

### Local immune response

Software for the deconvolution of immune cells in a tissue sample can be used to measure the local immune response to a tumor. Research in colorectal cancer demonstrated that the “immune contexture” of the tumor could be used to predict patient survival [244–247]. Tumors heavily infiltrated by anti-cancer immune cells are said to have “hot” tumor microenvironments (TMEs); tumors that are devoid of tumor infiltrating immune populations are considered “cold” [248]. Researchers have associated positive patient outcomes to effector T cells and mature dendritic cells [249] and poor outcomes to regulatory T cells [250, 251], myeloid-derived suppressor cells (MDSCs) [252], and tumor-associated macrophages (TAMs) [253, 254]. The poor outcomes may be related to immune infiltrate promoting metastasis [255].

The local immune response can be characterized not just by the cell types present but also by the states of those cells and their specific targets in the tumor. A recently proposed “cytolytic signature” attempts to capture cancer cell-killing activity of immune cells in the TME [256]. Other work has used MHC multimer staining [257, 258] and TCR-seq of tumor-infiltrating lymphocytes (TILs) [259–261] to enumerate the targets and magnitude of the local T cell response to cancer.

While the immune contexture can be deconvolved from RNA-seq of a bulk tumor sample, other measurements may provide more insight. Recent work has shown that the epigenetic profile of a cell, as measured by ATAC-seq, is more stable than RNA-seq of cells of the same type or in the same state [262]. Deconvolution algorithms that take ATAC-seq data as input will likely be more accurate than those that use RNA-seq. It is also likely that high-dimensional single-cell profiling technologies like mass cytometry or single-cell RNA-seq will provide a more detailed understanding of the immune contexture.

Immunohistochemistry (IHC) images capture the spatial distribution of immune cells, and new multiplex protocols allow staining of up to a few dozen antigens on a single slide [263, 264]. Beyond IHC, new techniques like histo-cytometry [265] and multiplexed ion beam imaging (MIBI) [266] make high dimensional quantification of cells possible while capturing their spatial distribution.

### Tumor-extrinsic immune escape

The TME is more than just tumor and immune cells: stromal cells, blood and lymphatic vessels, extracellular matrix, interstitial fluid, and microbiota all contribute to the immunogenicity of a tumor [267, 268]. The tumor vasculature is usually studied through the lens of preventing angiogenesis to starve the tumor of nutrients [269]. Blood vessels, however, also permit T cell trafficking to tumors and can be altered in cancer to facilitate immune escape [270, 271]. Tumors can also develop local lymphatics whose role in the immune response to cancer is still being elucidated [272].

Stromal cells modulate the immune contexture of the TME in multiple ways, including altering the contents of the tumor interstitial fluid or remodelling the extracellular matrix to exclude immune cells [273, 274]. They are also an important source of inhibitory checkpoint ligands [275]. Features of the extracellular environment that alter the immune contexture of the TME include ion [276], cytokine [277], and oxygen [278] concentrations, and the tumor microbiome [279, 280]. A particular area of recent focus is the competition for metabolites between clonally expanding tumor and T cells [281, 282]. While DNA-seq and RNA-seq data from tumor biopsies can be used to look for pathways known to modulate the TME, investigation of tumor-extrinsic immune escape is aided significantly by metabolomic data [283].

### Systemic immune response

Measurement of the tumor draining or sentinel lymph node [284] as well as profiling of the peripheral blood [285] can provide additional information about the immune response to cancer [286, 287]. In addition, the diversity of the peripheral adaptive immune receptor repertoire can be predictive of disease progression [288, 289], and probing the antigen specificity of the antibody [290] or T cell [291] repertoire can reveal an immune response to tumor antigens. PhIP-seq is a new technology that allows for low cost, high-throughput profiling of the antigen specificity of the antibody repertoire [292, 293].

Finally, it is well known that cancer incidence increases with age. Measures of aging in the immune system [294] could potentially be used to predict tumor/immune interactions and guide treatment decisions [295].

### Summary measures

Recently, it was suggested that the various measures of tumor/immune interactions should be integrated into a single “immunogram” [296, 297] or “immunophenotype” [217, 298, 299]. These summary measures and visualizations can be useful for identifying immune molecular subtypes, prognosis, and clinical decision support, or suggesting new combination therapies [300].

## Therapeutic interventions

The understanding gained from studying the interaction of the immune system and cancer can be used to design interventions that improve the immune response to cancer. While conventional cancer therapies have immunological effects [301, 302], here, three approaches to cancer immunotherapy will be discussed: therapeutic vaccination, checkpoint modulation, and adoptive cell transfer (ACT).

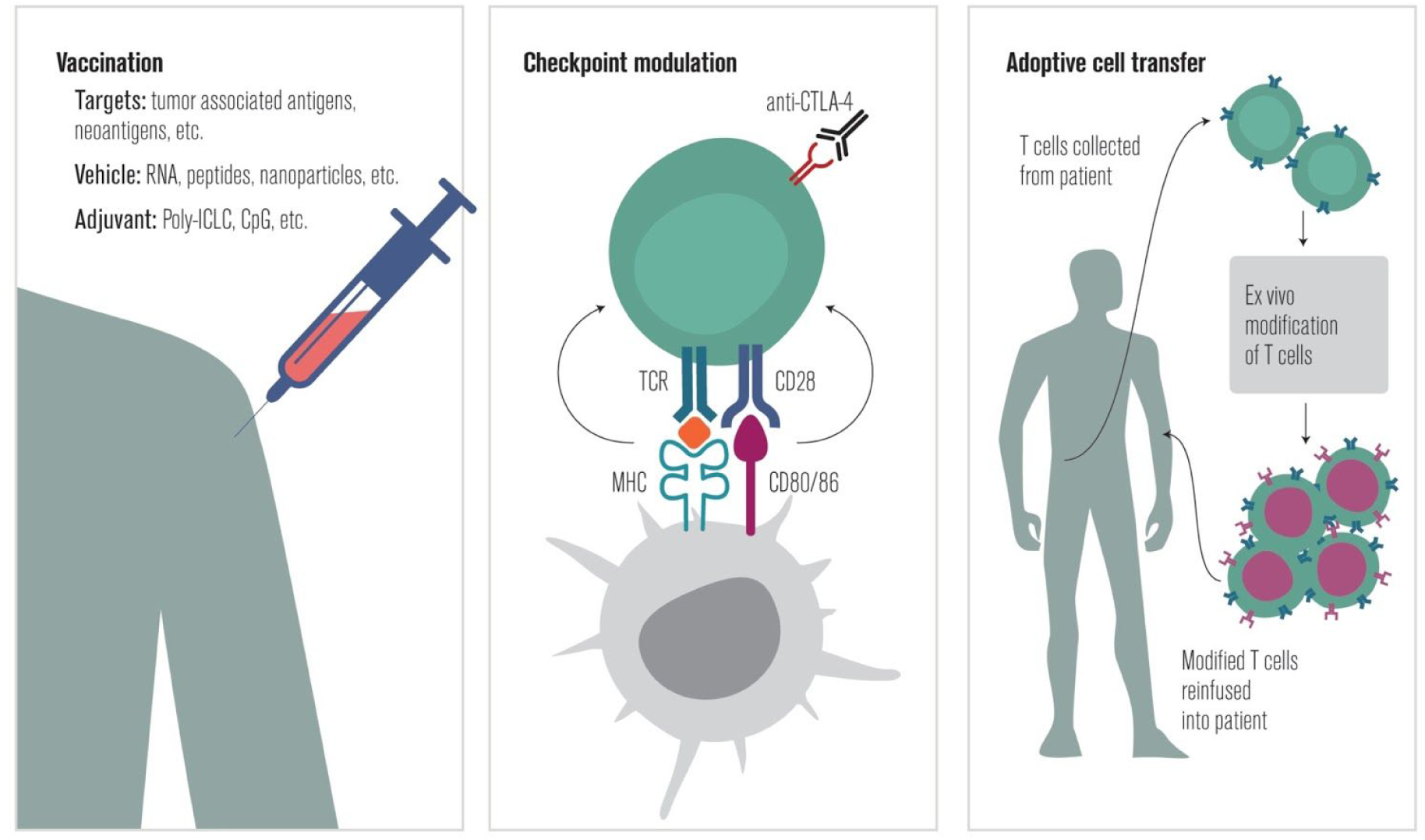

### Vaccination

In 2010, the FDA approved sipuleucel-T, a dendritic cell vaccine targeting the tumor antigen prostatic acid phosphatase, for use in prostate cancer [303]. Since 2011, CIMAvax-EGF, a vaccine targeting the growth factor receptor ligand EGF in lung cancer [304, 305], has been available in Cuba and is currently in clinical trials in the United States.

These approvals came after decades of development of many approaches to vaccination against established cancer [306–308] and inspired dozens of additional clinical trials [309–311].

Therapeutic cancer vaccines differ by target, delivery method, and adjuvants [312–315]. The major challenge in therapeutic cancer vaccination is not generating a tumor-directed immune response but rather overcoming immune evasion [316] and generating T cell memory [317]. Recently, several approaches to the personalization of therapeutic cancer vaccines have been proposed [318], including loading dendritic cells with material obtained from the tumor directly [319] or specifically targeting neoantigens [320–322].

The efficacy of neoantigen vaccines was demonstrated in mouse models in 2014 [258], and the first publication of results in human trials was in 2015 [323]. The neoantigens targeted by these vaccines are often identified by neoepitope prediction from sequencing or mass spectrometry data [324], though ranking factors may need to be tuned to this task: what makes a good endogenous neoantigen may not be identical to what makes a good target for neoantigen vaccination.

One surprising finding from the analysis of the first neoantigen vaccine responses was that class II epitopes are important for vaccine effectiveness even when vaccine epitopes are predicted based on MHC I prediction algorithms [325, 326]. An important concern with neoantigen vaccines is that some tumors may not have enough neoantigen-generating mutations to target [327].

In addition to target selection, bioinformatics can be used to optimize vaccine delivery and adjuvant composition. Prediction of how the breadth, intensity, and duration of the immune response to vaccination relate to antigen load, schedule, and injection site is an active area of research [328–331].

### Checkpoint modulation

The relatively recent success of checkpoint blockade therapies in melanoma [332], followed by other malignancies [333–336], has motivated a resurgence of research into immune checkpoint receptors and their ligands [337, 338]. The FDA has approved antibodies targeting CTLA-4 and PD-1/PD-L1, and new immune checkpoint targets are the subject of active investigation [339–343].

Because pembrolizumab was first approved in conjunction with a companion diagnostic, the IHC 22C3 pharmDx test [344, 345], there has been an intense focus on the association of PD-L1 level with response to blockade of the PD-1/PD-L1 axis [346]. Using PD-L1 level as a biomarker is difficult: PD-L1 is expressed by many cell types, its expression in those cells varies over time [275], its function varies depending on the cell type on which it is expressed [347], there are multiple causes of its presence or absence [348, 349], and it appears to interact with other biomarkers [350]. Nevertheless, work continues in order to understand the role of PD-L1 expression in response to other therapies [351] and in other indications [352]. The Blueprint PD-L1 IHC Assay Comparison Project recently reported their Phase 1 results comparing four assays for measuring PD-L1 [353].

Beyond immune checkpoint receptor or ligand expression, commonly proposed biomarkers for response to checkpoint blockade include mutation burden [354–356], mismatch-repair status [357], and loss of the interferon response pathway [358–361]. The composition of the gut microbiome [362, 363] and overall immune system state (as inferred from peripheral blood samples) [364–366] have also been implicated.

A major challenge for the field is to formulate a mechanistic model of checkpoint blockade that will allow predictive biomarkers that associate with response to intervention to be separated from prognostic biomarkers that associate with disease progression unrelated to intervention [367–369]. Lessons about which cells respond to checkpoint blockade from chronic infection [262, 370] may be important in the construction of the mechanistic models.

Additional challenges include finding biomarkers that predict adverse events [371–373], developing on-treatment biomarkers that predict response to therapy early in the course of treatment [374, 375], and the interpretation of predictive models to elucidate mechanisms of resistance [376] and suggest new combination therapies [377–379].

### Adoptive cell transfer

In 1988, it was reported that TILs from melanoma, when expanded *in vitro* and reinfused into the host, could cause tumor regression [380]. In 2002, host lymphodepletion prior to adoptive cell transfer (ACT) was used to improve response rates [381]. For many tumors, however, TILs are not available, and T cells specific for tumor antigens are created *in vitro* through genome editing. Autologous lymphocytes from the peripheral blood edited to express either a cancer antigen-specific TCR [382] or a cell-type specific chimeric antigen receptor (CAR) [383] have been used to treat cancer in humans. For more detailed reviews of ACT, see [384] and [385].

One primary use of bioinformatics in ACT is to identify new targets for cell therapy [386–388]. Similar techniques to those used to discover targets for vaccination are employed in this domain. Bioinformatics can also examine how the targets evolve in response to ACT [389, 390].

To determine the optimal cell product, detailed measurements are performed on cells prior to and after transfer. An increase in the number of T cell targets [391] and the persistence of anti-tumor T cells [392] are post-transfer features associated with positive outcomes. Evolving technologies enable more detailed tracking of the fate of adoptively transferred cells *in vivo* [393], including with PET [394–396].

Another use of bioinformatics in ACT is to discover the expansion and enrichment protocol that generates the optimal cell product for reinfusion. Cytokines [397], metabolites [398], and small molecules [399] have all been shown to impact therapeutic efficacy. Artificial antigen presenting cells (aAPCs) can also be used [400].

Finally, informatics can help determine how to best prepare the patient for ACT. Total body irradiation and chemotherapy are commonly used to deplete lymphocytes in the host prior to ACT; it is not yet clear whether intensive myeloablative lymphodepletion, which requires hematopoietic stem cell transplantation (HSCT) to reconstitute the host immune system, is more effective than transient, nonmyeloablative lymphodepletion [401–403].

## Conclusion

Informatics techniques to analyze tumors, the tumor microenvironment, and systemic immunity have evolved alongside the increased use of immunomodulatory therapies in cancer. The rate of progress has been dizzying: the first published use of NGS to predict neoantigens involved spreadsheets and manual curation [211]. Just four years later, the Schreiber lab and collaborators had streamlined the process into a pipeline [404]. Immune deconvolution techniques have enriched the types of data that can be extracted from existing RNA-seq or microarray data, giving a more complete picture of the tumor and its immune microenvironment. This deepened understanding is being brought to the next level by techniques such as single cell RNA-seq and multiplex ion beam imaging (MIBI).

However, challenges abound. First, the determination of sample quality and tumor proportion (in the case of human tumor samples) is uncertain and serves as an unstable foundation for the intricate and often elegant work of translational bioinformatics. Second, as demonstrated by the array of software packages that can be used at each stage of the analytic process, there is no single accepted or superior method with which to perform each step. Consequently, research groups working with distinct methods may generate substantially different results and conclusions from the same dataset. Third, the application of statistical methods to the “multi-omics” data being generated poses a substantial challenge, especially given that many translational studies feature small sample sizes and many studied variables: a perfect storm for false discovery, a risk of which the field must be critically aware. Finally, as preclinical data has demonstrated, metabolomics likely play an important role in tumor immunity, but quality data is difficult to collect when using human samples.

As bioinformatics techniques continue to improve, they are likely to play an ever-growing role in the triage of patients to off-the-shelf therapies, as in the case of mutation load with checkpoint blockade agents, or be used to determine what therapy is given, as in the case of vaccination or ACT.

## Acknowledgements

The authors would like to thank Arman Aksoy for his input to this review.

